# Bulk transcriptomic subtypes in high-grade serous ovarian carcinoma (HGSC) are readouts of tumor cellular composition

**DOI:** 10.64898/2026.04.16.719000

**Authors:** Stephanie Tanis, Manoel Lixandrão, Adriana Ivich, Laurie Grieshober, Katherine A. Lawson-Michod, Lindsay J. Collin, Lauren C. Peres, Lucas A. Salas, Jeffrey R. Marks, Benjamin G. Bitler, Casey S. Greene, Joellen M. Schildkraut, Jennifer Anne Doherty, Natalie R. Davidson

**Affiliations:** Department of Obstetrics and Gynecology, Division of Reproductive Sciences, University of Colorado Anschutz, Aurora, CO 80045, USA; Department of Biomedical Informatics, University of Colorado Anschutz, Aurora, CO 80045, USA; Huntsman Cancer Institute, University of Utah, Salt Lake City, UT 84112, USA; Department of Population Health Sciences, University of Utah, Salt Lake City, UT 84108, USA; Division of Public Health Sciences, Fred Hutchinson Cancer Center, Seattle, WA 98109, USA; Department of Epidemiology, University of Washington, Seattle, WA 98109, USA; Department of Epidemiology, Rollins School of Public Health, Emory University, Atlanta, GA, 30322, USA; Department of Cancer Epidemiology, Moffitt Cancer Center, Tampa, FL 33612, USA; Department of Epidemiology, Geisel School of Medicine, Dartmouth, Lebanon, NH 03784, USA; Department of Surgery, Duke University School of Medicine, Durham, NC 27707, USA

## Abstract

High-grade serous ovarian carcinoma (HGSC) is classified into four transcriptomic subtypes that are reproducible across cohorts but have not informed treatment. These subtypes are associated with stromal and immune content, yet they are assigned from bulk expression, from which composition is also estimated. We tested whether composition alone determines subtype. In tumors constructed from single cells with epithelial cells held constant, varying a single cell type within the range observed in primary tumors reassigned the subtype, and known composition distinguished all four subtypes. Varying epithelial cell identity at fixed composition displaced tumors approximately half as far in expression space. Across 2,161 primary tumors from ten cohorts, composition structured the leading axes of expression and predicted subtype in held-out cohorts (ROC-AUC 0.817 to 0.963). The consequence is prognostic. Adjustment for composition largely attenuated subtype associations with survival, whereas fibroblast content remained prognostic. Subtype-associated biology and prognosis therefore largely reflect the tumor microenvironment.

## INTRODUCTION

Bulk transcriptomic profiling is the most widely used approach for characterizing molecular heterogeneity across cancer types (*1*, *2*), and in ovarian high-grade serous carcinoma (HGSC) it has produced a framework the field has used to organize that heterogeneity for fifteen years. Ovarian cancer is the second leading cause of gynecologic cancer death, and HGSC, as its most common histotype, accounts for the majority of those deaths (*3–5*). Despite near-universal *TP53* mutation and broadly similar patterns of chromosomal instability, HGSC tumors differ substantially in gene expression, immune and stromal infiltration, and clinical outcome (*2*, *6–9*). Four transcriptomic subtypes have organized that heterogeneity since they were defined: C1.MES (MES; mesenchymal), C2.IMM (IMR; immunoreactive), C5.PRO (PRO; proliferative), and C4.DIF (DIF; differentiated) (*2*). They are reproducible across independent cohorts and profiling platforms and are used routinely to interpret tumor biology and to contextualize molecular and clinical associations (*10–14*), but they have not produced stratification that changes treatment (*15*). Their role has been largely interpretive, which makes it important to know what each label measures.

The subtypes are assigned from bulk profiles, and a bulk profile is a weighted average over epithelial, stromal, immune, and endothelial cells, such that the patterns recovered from it reflect the relative abundance of those compartments as readily as the transcriptional state of the cells within them (*16–18*). The subtypes have been linked to composition accordingly. Single-cell and integrative analyses associate MES and IMR expression with fibroblast and immune infiltration respectively (*19–26*), and pan-cancer work shows that tumor purity shifts bulk expression profiles and subtype assignment (*17*, *27*). Mixing microdissected HGSC epithelium with its paired stroma showed that adding 10-30% stroma reassigns subtype, and that several published prognostic signatures lose their survival association once stromal content is accounted for (*28*). In colorectal cancer, the mesenchymal and immune consensus molecular subtypes were similarly traced to stromal and immune content (*29*, *30*), a conclusion later confirmed by assigning subtypes to individual cells (*31*). These studies observed composition and subtype as they co-occur in existing tumors, or varied a single stromal fraction, often with composition estimated by deconvolution from the same expression matrix that produced the label (*32*). They establish that stroma can move the call, but not which cell types do so within the range primary tumors span, how much of each label composition explains, or whether the dependence extends beyond MES and IMR to DIF and PRO.

Answering these questions requires tumors whose composition is specified by construction rather than estimated from their expression, with epithelial cell state held constant as composition varies, and with that variation kept within the range where the labels are used. Subtype frameworks were derived from bulk profiles of surgically resected tumors selected for tumor content (*2*, *33*) and are applied to profiles assembled the same way, so the compositional range those tumors span is the range in which the labels are defined. Both conditions became achievable once single-cell atlases of HGSC grew large enough to reconstitute tumors from real cells (*34*, *35*).

Here we set cellular composition directly, using a published single-cell HGSC atlas (*34*) to construct pseudobulk tumors in which composition is specified while epithelial transcriptional programs are held constant. We then invert the design and vary which patient’s epithelial cells, or which anatomical site within a patient, populates the epithelial compartment at fixed composition. We ask whether the resulting dependencies hold in 2,161 primary HGSC transcriptomes from ten cohorts (*2*, *20*, *36–42*), profiled in the same way as those from which the subtypes were derived, with composition inferred by deconvolution (*18*, *43*). In those tumors, we locate the dependencies within the global structure of bulk expression, test whether composition alone recovers subtype labels in cohorts held out entirely, and test whether subtype retains any association with overall survival once composition is measured. Changing composition changes the subtype call, and when composition is known by construction it separates all four subtypes, including DIF and PRO. Epithelial cell identity moves tumors about half as far, but enough to change the call for tumors near a class boundary. A subtype label therefore divides a continuous compositional plane into four regions, and what it reports can be measured directly from data that already exist.

## RESULTS

### Subtype identity tracks the composition of the mixture

Subtype labels are assigned from mixtures, so we first measured the effect of the mixture directly. From a single-cell HGSC cohort of 41 patients (*34*) we drew a random subset of 500,000 droplets, yielding 337,765 annotated cells across seven anatomical supersites and 137 patient-site combinations (Fig. S1). Each tumor was reconstituted as a pseudobulk profile from its own annotated cells, giving composition that is counted rather than inferred, and then rebuilt from its epithelial cells alone (Fig. 1A, Table S1). Thirty-nine patients had at least 500 total and 200 epithelial cells and therefore entered both builds (Fig. S1D). These tumors differed from one another in composition by up to 25-fold, spanning 3-fold in epithelial content, 11-fold in fibroblast and 25-fold in mast cell across the 39 patients (Fig. 1B), and occupied a wider and less epithelial compositional range than the bulk cohorts these labels are usually applied to (Fig. S4A).

**Figure 1.**
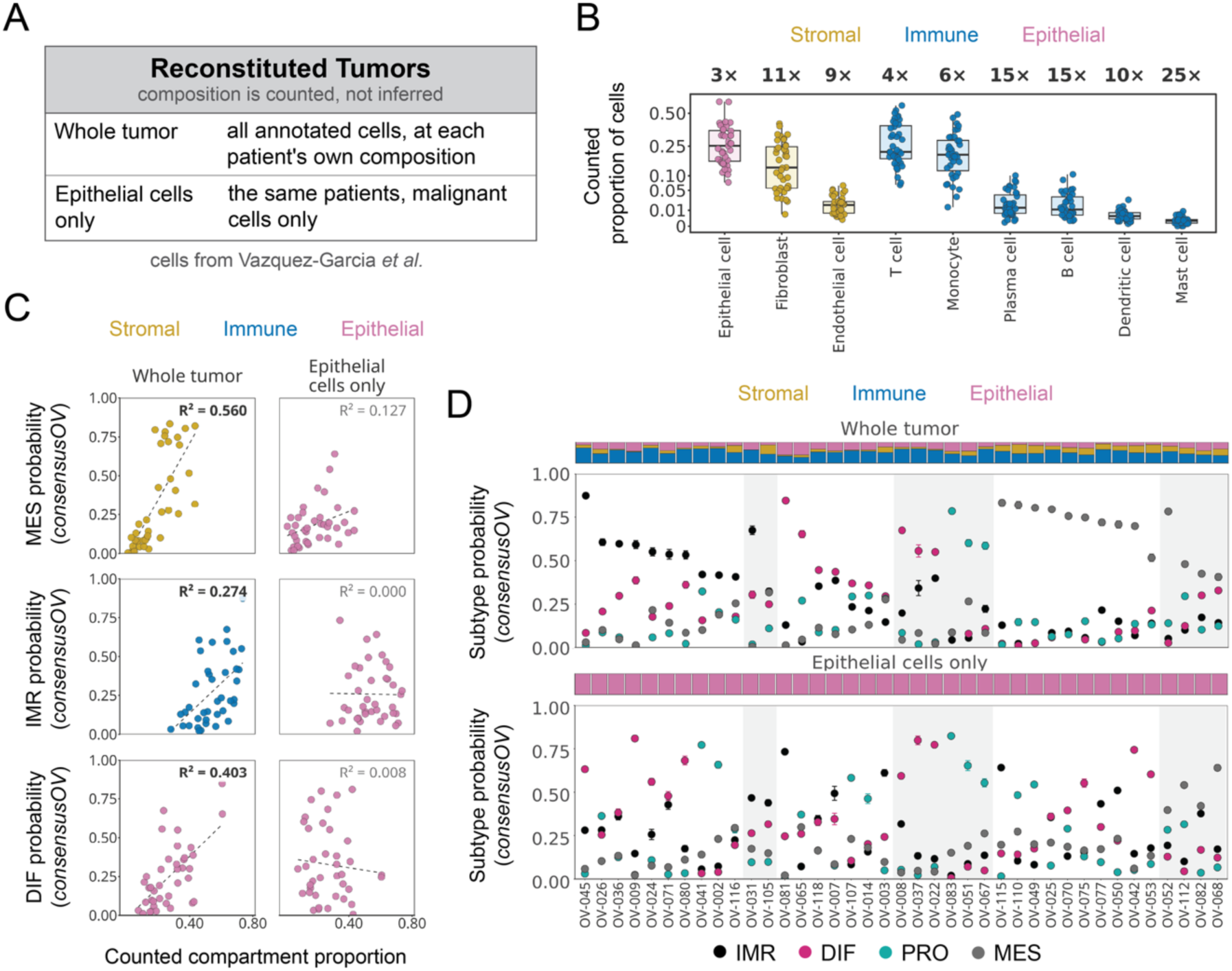
Subtype assignment in tumors reconstituted from annotated single cells. **(A)** The two builds. Each of 39 patients was reconstituted as a pseudobulk profile from all of that patient’s annotated cells at the patient’s own composition, and again from that patient’s epithelial cells alone. Cells are from Vázquez-García et al. (34). Composition in these profiles is counted from the annotations rather than inferred. **(B)** Counted proportion of each compartment across the 39 patients, on a square-root scale. Boxes show the median and interquartile range with whiskers to 1.5 times the interquartile range; each point is one patient. Values above each compartment are the fold range between the 10th and 90th percentiles. Compartments are grouped as epithelial, stromal and immune. **(C)** Counted compartment proportion against consensusOV subtype probability, for the whole-tumor builds (left) and the epithelial-only builds (right). Rows are the subtype most strongly associated with each compartment in the whole-tumor builds. Compartments are collapsed to stromal (fibroblast and endothelial cell), immune (all remaining) and epithelial; in the epithelial-only builds the epithelial proportion is 1 by construction and the x axis is the whole-tumor value. Dashed lines are ordinary least squares fits with R² annotated. Associations with R² > 0.195 are shown in bold. All 36 compartment and subtype pairings are given in Fig. S2. **(D)** consensusOV subtype probabilities for each patient, from the whole-tumor builds (upper) and the epithelial-only builds (lower). Points are means across 10 replicate pseudobulk profiles with bars spanning the 95% confidence interval. The strip above each panel is that patient’s counted composition in the three compartments. Patients are ordered by their whole-tumor call, and shaded bands mark the 12 patients whose epithelial-only call matched it.

Counting composition rather than inferring it makes the association between composition and subtype directly measurable, without the circularity of estimating both from one matrix. Across the 39 patients, MES probability increased with counted stromal content (R² = 0.560), DIF probability with epithelial content (R² = 0.403), and IMR probability with immune content (R² = 0.274) (Fig. 1C, left). At cell type resolution, six of the 36 compartment and subtype pairs were significant after correction, all involving fibroblast, epithelial cell or T cell (Fig. S2, Table S2). The stromal association was carried by fibroblast and the immune association by T cell content (R² = 0.208), whereas monocyte content was unrelated to IMR probability (R² = 0.014).

Rebuilding the same tumors from epithelial cells alone largely removed the associations of MES with stromal content, DIF with epithelial content, and IMR with immune content. Against the same counted proportions, MES probability retained a weaker association (R² = 0.127), and IMR and DIF probabilities showed none (R² = 0.000 and 0.008) (Fig. 1C, right). The relationships in the whole-tumor builds were therefore carried by the compartments removed, not by the epithelial cells that remained, which were identical between the two builds. The maximum-probability call changed in 27 of 39 patients, and 12 retained the call their whole-tumor profile received (Fig. 1D). Call margins were comparable between the two builds (median 0.335 and 0.292, paired Wilcoxon p = 0.34), so the change is a reassignment rather than a loss of confidence (Fig. S3). Removing those cells alters composition and the epithelial expression average together, so what this establishes is that the label depends on what else is in the mixture, not which of the two moves it. Separating them requires composition to be set rather than observed.

### Composition reassigns subtypes within biologically plausible tumor compositions

To vary composition independently of epithelial cell state, we constructed pseudobulk mixtures from the same annotated cells in which the proportion of each of the nine major compartments is set by design and the epithelial cells entering every mixture are held fixed. Four designs were used, with experimental control decreasing at each step: one compartment titrated against a fixed background, epithelial donor varied at fixed composition, anatomical site varied at fixed composition and donor, and all compartments varied at once (Fig. 2A).

**Figure 2.**
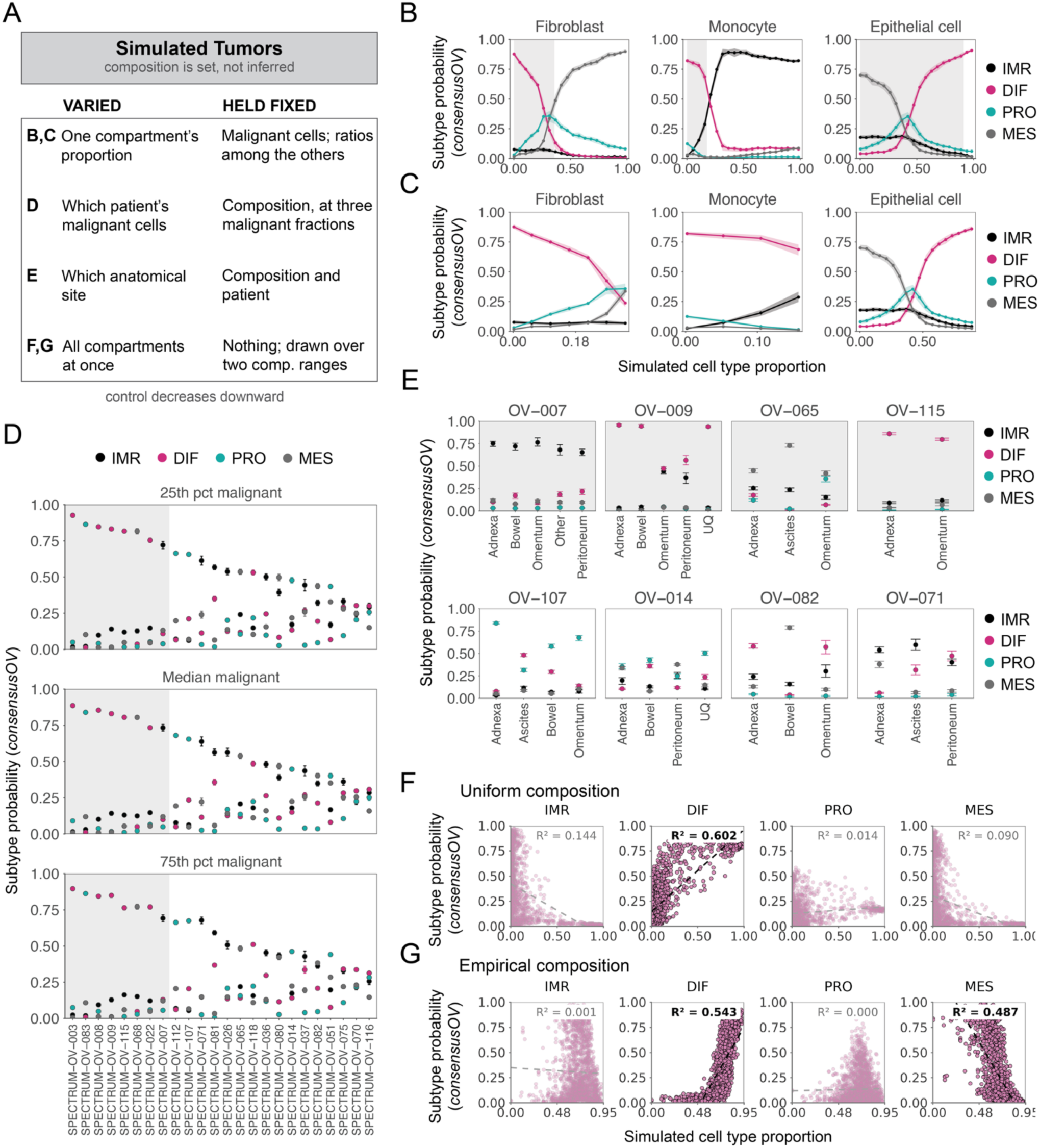
Subtype assignment when composition is set by design. **(A)** The four designs, with what varies and what is held fixed in each. Experimental control decreases downward. **(B)** One compartment stepped from 0 to 1 in 20 steps while the ratios among the remaining compartments are held fixed, with 30 replicate mixtures per step. Points are means across replicates with ribbons spanning the interquartile range across replicates. Shading marks the compositional range primary tumors occupy, defined as zero to the 99th percentile of that compartment across the 2,161 primary profiles. All nine compartments are given in Fig. S5A. **(C)** The same titrations restricted to that range. **(D)** Composition held at the primary-cohort median while the patient supplying the epithelial compartment is varied, at three epithelial fractions: the 25th percentile, median and 75th percentile of the primary cohorts (0.633, 0.712 and 0.783). Twenty-four patients contributed at least 1,500 epithelial cells and entered this design. Points are means across replicates with bars spanning the 95% confidence interval. Donors are ordered by their highest subtype probability at the 25th percentile fraction, and that order is held across all three panels. Shading marks the 8 donors whose call was the same at all three fractions with a margin over the runner-up of at least 0.5. **(E)** Composition and patient held fixed while the anatomical site supplying the epithelial cells is varied, for eight of the 23 patients with two or more separately sampled sites, four whose call was the same at every site (shaded) and four whose call switched. Points are means across replicate mixtures with bars spanning the interquartile range. All 23 are given in Fig. S6B. **(F, G)** All nine compartments varied simultaneously, 1,900 mixtures per scheme, under compositions drawn uniformly over the simplex (F) and compositions resampled from the inferred compositions of the primary cohorts (G). Each point is one mixture; epithelial proportion is on the x axis in every panel. Dashed lines are ordinary least squares fits with R² annotated. Associations with R² > 0.195 are shown in bold. All nine compartments are given in Fig. S7.

Titrating one compartment while holding the rest fixed isolated that compartment’s effect on the call. Increasing fibroblast content moved the maximum-probability call from DIF to PRO at 0.158 and from PRO to MES at 0.211, increasing monocyte content moved it from DIF to IMR at 0.211, and increasing epithelial cell content moved it from MES to PRO at 0.632 and from PRO to DIF at 0.684 (Fig. 2B). Proportions sum to one, so these reflect compositional trade-offs rather than independent effects of single compartments.

Restricting the simulation to the compositional range primary tumors occupy retained the fibroblast and epithelial transitions inside the observed window, while the monocyte transition fell just outside it, at 0.211 against an observed maximum of 0.177 (Fig. 2C). Endothelial cells crossed at the edge of the window, at 0.263 against an observed maximum of 0.275, and the remaining compartments only at proportions 5 to 74 times higher than any primary tumor reached (Fig. S5A, Table S3). To test whether these positions depend on the background the titration was anchored at, we repeated it with the epithelial fraction set at the 25th, 50th and 75th percentiles of primary tumors and with a second single-cell reference (GSE217517). Fibroblast and monocyte crossovers moved by at most 0.053 and epithelial crossovers by at most 0.105 (Fig. S5A). Deconvolution recovered fibroblast, epithelial, and monocyte proportions with fitted slopes of 0.936 to 1.051 against known proportions, and no compartment fell below 0.70, a bias smaller than the 2- to 16-fold differences between reconstituted and primary medians (Fig. S4B to D).

Varying the epithelial donor while holding composition fixed isolated the effect of epithelial identity on the call. Changing which patient’s epithelial cells filled that compartment moved subtype probability across most of its range (Fig. 2D). The design requires enough epithelial cells from one patient to fill the compartment at the fixed epithelial fraction, so it draws on the 24 patients contributing at least 1,500 epithelial cells. Donor rank order was preserved at three epithelial fractions (Fig. 2D), and 22 of 24 donors retained their call when each donor’s pool was left at its natural depth rather than equalized (Spearman ρ = 0.989 for IMR, 0.972 for DIF, 0.980 for PRO and 0.988 for MES; Fig. S6A), so the effect reflects donor identity rather than sampling depth. Varying only the anatomical site within a patient, at fixed composition and donor, draws on the 23 patients with two or more separately sampled sites. The call was stable across sites in 15 of these patients and switched in the remaining 8, most often between DIF and IMR (Fig. 2E, Fig. S6B).

Releasing all nine compartments at once removes the fixed background the titration relied on, so what varied now depended entirely on the compositions the mixtures were drawn from. Under compositions drawn uniformly at random, epithelial fraction explained most of the variance in DIF probability (R² = 0.604) and little in the others (IMR 0.145; MES 0.094; PRO 0.016) (Fig. 2F). Under compositions resampled from primary tumors, the same axis explained substantial variance in DIF (R² = 0.537) and MES (R² = 0.515) and almost none in IMR (R² = 0.001) or PRO (R² = 0.004) (Fig. 2G). At cell type resolution, uniform sampling produced associations of epithelial cell with DIF (R² = 0.604), endothelial cell with PRO (0.522), fibroblast with MES (0.496) and monocyte with IMR (0.284), while sampling from primary tumors produced the strongest associations for fibroblast with MES (0.709), epithelial cell with DIF (0.537) and MES (0.515), and monocyte with IMR (0.512) (Fig. S7). Sampling from primary tumors shifted the dependence on epithelial fraction from DIF alone to DIF and MES.

### Compositional dependencies in 2,161 primary tumors

We assembled 2,161 primary, treatment-naive HGSC transcriptomes across ten cohorts, 898 by RNA-sequencing and 1,263 by microarray (Fig. 3A, Table S4), and inferred proportions for the same nine compartments using InstaPrism (43). TCGA contributes both platforms, and 226 of its tumors were profiled on both. Both profiles are retained in these descriptive analyses, and the duplicate array profiles are removed wherever tumors are treated as independent observations (Methods). These cohorts were assembled under tumor-content criteria (*2*) and are correspondingly epithelium-rich. Median epithelial content ranged from 0.633 to 0.793 across cohorts, and epithelial cells were the largest compartment in all ten (Fig. S8, Table S5). Inferred proportions and subtype probabilities are both derived from the same expression matrix and are not independent measurements, so what follows asks whether the dependencies established by design appear at scale, and how much of the label they explain.

**Figure 3.**
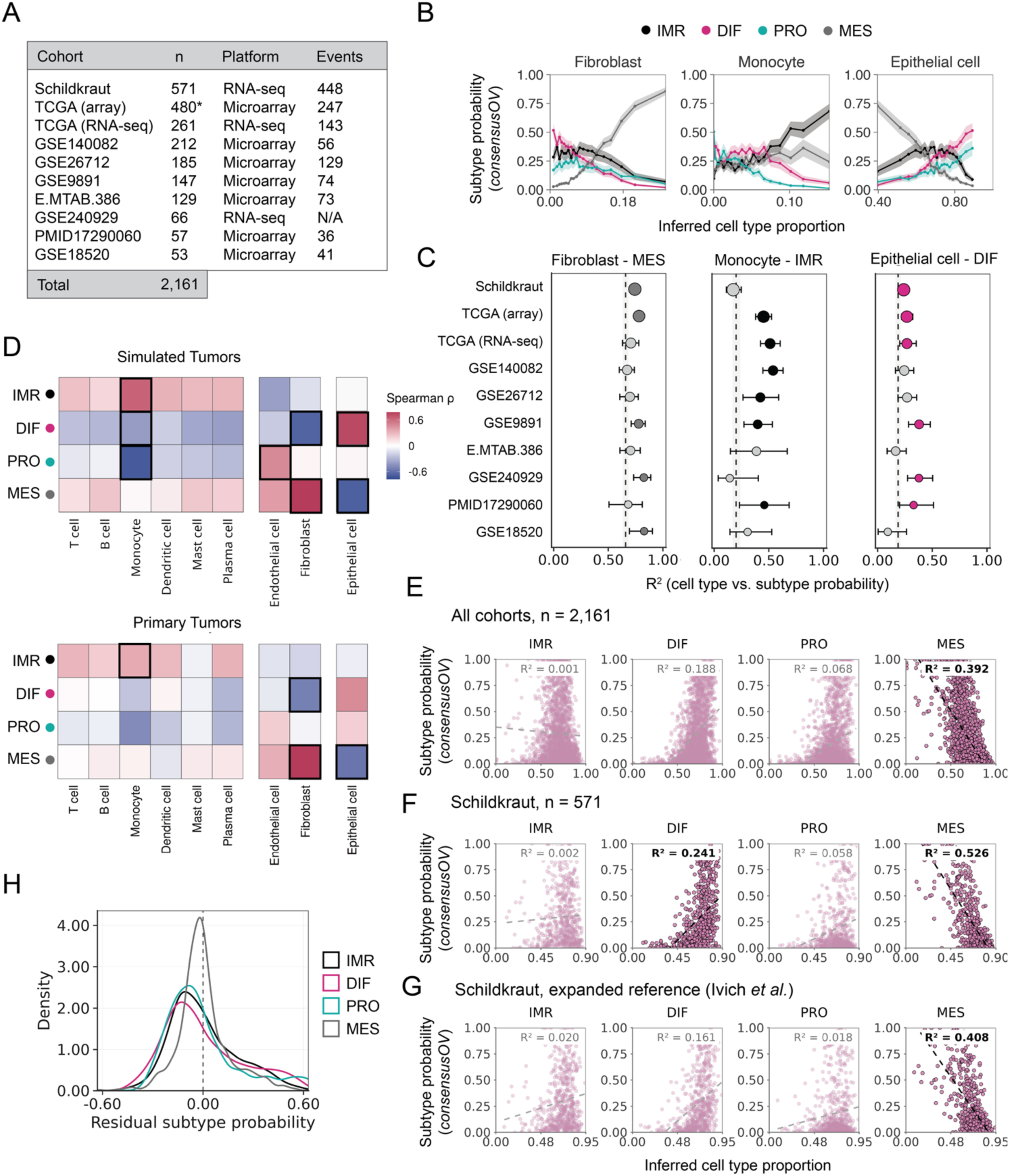
Compositional dependencies in 2,161 primary tumors. **(A)** The ten cohorts, with sample count, profiling platform and number of deaths. Asterisk marks the TCGA microarray cohort, of which 226 patients are also present in the TCGA RNA-seq cohort. N/A indicates no survival annotation. Full cohort characteristics, including follow-up and the clinical covariates available in each, are given in Table S4. **(B)** Inferred compartment proportion against consensusOV subtype probability across all 2,161 tumors, binned along the x axis with ribbons spanning the 95% confidence interval of the mean within each bin. Axis ranges are the observed range of each compartment; the distribution of each compartment within each cohort is given in Fig. S8A. **(C)** Variance in subtype probability explained by compartment proportion (R²) within each cohort, for the three pairings in B. Bars span the 95% confidence interval, and point size scales with cohort size. The dashed line and shaded band are the pooled R² and its 95% confidence interval across all 2,161 tumors. Points are colored by subtype where the cohort interval lies above the pooled value and grey where it includes it. **(D)** Spearman correlation between every compartment and every subtype probability, in the simulated tumors under empirical composition sampling (upper) and the primary tumors (lower). Cells are outlined where R² > 0.195. Adjusted p values for every pairing are in Table S2. **(E-G)** Epithelial cell proportion against subtype probability, in all cohorts pooled (E), in Schildkraut alone (F), and in Schildkraut deconvolved against the expanded fourteen-population reference (44) (G). Dashed lines are ordinary least squares fits with R² annotated, bold where R² > 0.195. All 36 compartment and subtype pairings are given in Fig. S9 for E and F, and all 56 for the expanded reference in Fig. S10. **(H)** Residuals from a model of subtype probability on all nine inferred proportions, across the 2,161 tumors. The model explained 0.672 of the variance in MES, 0.503 in IMR, 0.254 in DIF and 0.247 in PRO, with residual standard deviations of 0.161, 0.205, 0.238 and 0.230.

To compare the two settings on the same terms, we plotted subtype probability against inferred proportion over the compositional range these cohorts occupy. The curves followed those obtained by simulation (Fig. 3B; compare Fig. 2C). MES probability rose with fibroblast content and overtook the other subtypes at 0.058, IMR rose with monocyte content, and DIF rose with epithelial cell content as MES fell. Within each cohort, fibroblast content explained 0.668 to 0.822 of the variance in MES probability, monocyte content 0.149 to 0.547 of IMR, and epithelial content 0.098 to 0.382 of DIF (Fig. 3C). The fibroblast relationship was the strongest of the three in every cohort. At cell type resolution, four of the 36 compartment and subtype pairs exceeded R² = 0.195, involving fibroblast, epithelial cell and monocyte content (Fig. S9A).

Comparing the associations that exceeded R² = 0.195 in each setting identified where the two agreed (Fig. 3D). All four in primary tumors, fibroblast with MES and DIF, epithelial cell with MES and monocyte with IMR, also exceeded it in simulated tumors resampled from primary compositions. Four exceeded it only in simulated tumors: epithelial cell with DIF (R² = 0.537 against 0.185 in primary tumors), monocyte with PRO (0.347 against 0.151) and DIF (0.216 against 0.052), and endothelial cell with PRO (0.223 against 0.006) (Fig. S7B, Fig. S9A). In primary tumors, where the remaining compartments vary alongside the one plotted rather than being held fixed, the fibroblast transition fell at 0.058 rather than 0.211 and the monocyte transition at 0.003 rather than 0.211. The epithelial transition matched closely (0.644 against 0.632).

Quantifying the dependency requires asking how much of each subtype probability composition explained. Across the pooled cohorts, epithelial fraction alone explained substantial variance in MES (R² = 0.390) and DIF (R² = 0.185) but essentially none in IMR (R² = 0.002) or PRO (R² = 0.068) (Fig. 3E). All nine compartments together explained 0.672 of the variance in MES, 0.503 in IMR, 0.254 in DIF and 0.247 in PRO. Residuals from this model were tightest for MES (SD = 0.161) and widest for DIF (SD = 0.238), with IMR and PRO intermediate (SD = 0.205 and 0.230) (Fig. 3H).

Two features of the cohorts could have produced or inflated these values: pooling across platforms, and the resolution of the deconvolution reference. Restricting to a single cohort on a single platform strengthened rather than weakened the relationship (Schildkraut, n = 571; MES R² = 0.516; DIF R² = 0.231; Fig. 3F, Fig. S9B). Deconvolving the same cohort against an independent reference (*44*), which resolves fourteen cell populations rather than nine but remains a coarse partition of HGSC tissue, weakened it but retained most of it (MES R² = 0.400; DIF R² = 0.153; Fig. 3G, Fig. S10). The finer reference redistributed the immune associations. NK cell and macrophage content showed the strongest associations with IMR probability (R² = 0.185 and 0.165), both below the 0.195 threshold, while monocyte content showed none (R² = 0.000) (Fig. S10).

### Cellular composition structures the leading axes of bulk expression

To locate this dependency within the global structure of bulk expression rather than in pairwise associations, we performed principal component (PC) analysis on 5,000 high-variance genes shared across all cohorts. Each of the 226 TCGA patients profiled on both platforms was represented once, by its RNA-seq profile, leaving 1,935 unique tumors.

The first three components explained comparable fractions of variance (PC1 22.6%; PC2 17.9%; PC3 15.5%), and the four subtypes occupied distinct but overlapping regions of the PC1-PC2 plane (Fig. 4A). Subtype identity explained 55.1% of the variance along PC1, 20.2% along PC2 and 26.5% along PC3 (Fig. 4B). Each of the three leading components corresponded to a compositional contrast (Fig. S12). PC1 correlated positively with epithelial cell content (ρ = 0.624) and negatively with monocyte, fibroblast and plasma cell content (ρ = −0.524, −0.524 and −0.370). PC2 contrasted epithelial against fibroblast and endothelial cell content (ρ = 0.376, −0.464 and −0.374), and PC3 carried a further fibroblast against epithelial cell contrast (ρ = −0.487 and 0.356) (Fig. 4B, D). Each of the first five components correlated with at least one compartment at |ρ| > 0.24.

**Figure 4.**
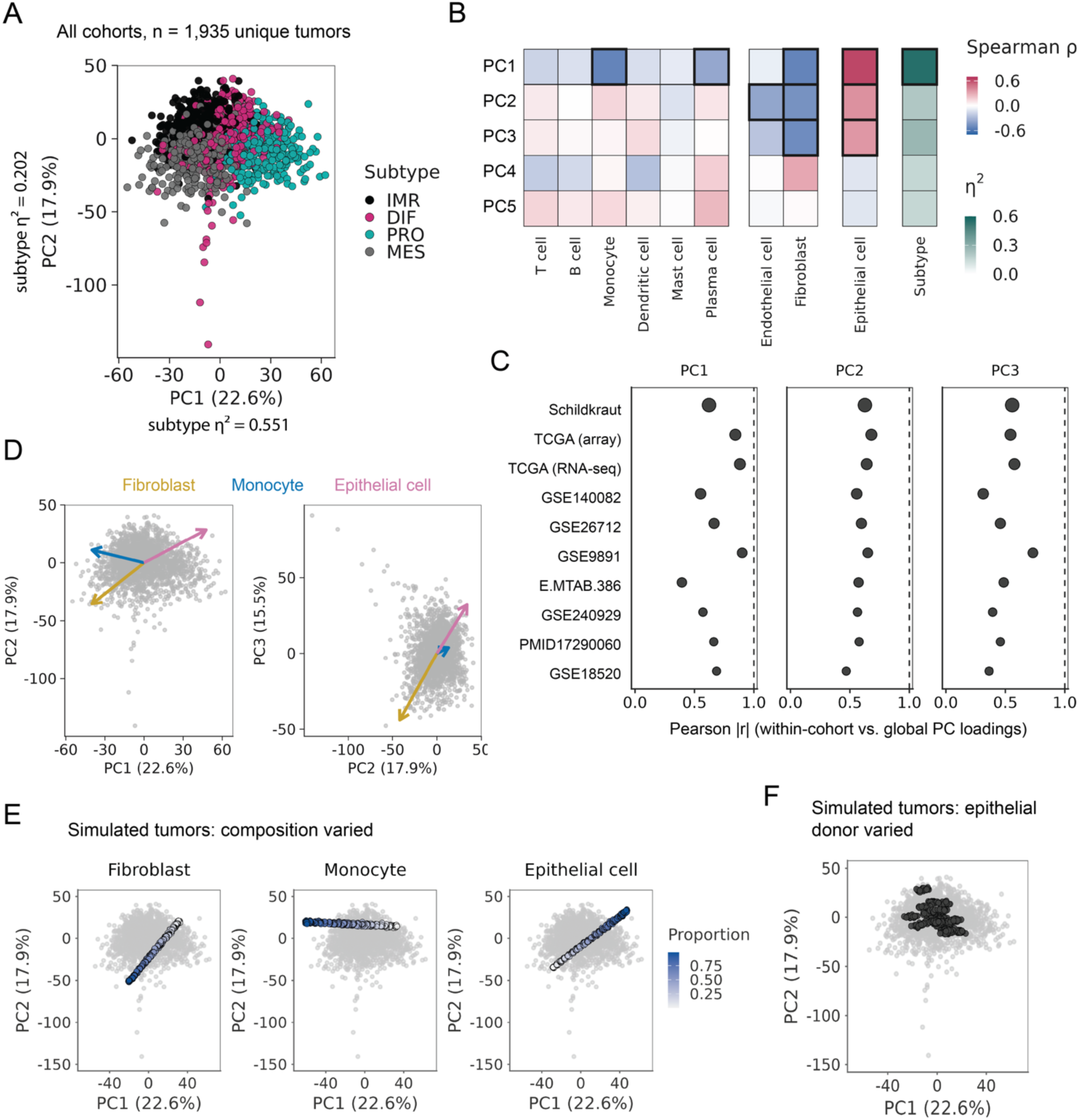
Cellular composition and the leading axes of bulk expression. **(A)** Principal component analysis of 1,935 unique tumors on 5,000 shared genes, colored by consensusOV call. Axis labels give the variance explained. Subtype identity explained 55.1% of the variance along PC1 and 20.2% along PC2. The same tumors colored by cohort and by platform, with the variance each account for, are given in Fig. S11A-C. **(B)** Spearman correlation between each of the first five components and each inferred compartment proportion, with eta squared for subtype in the rightmost column. Cells with |ρ| > 0.44, equivalent to R² > 0.195, are outlined. **(C)** Absolute Pearson correlation between each cohort’s own loadings, from a principal component analysis fitted within that cohort, and the global loadings. Each global component was compared against every within-cohort component and the strongest absolute match retained; the dashed line marks perfect agreement. Fig. S11D shows the same values against the same-index component. **(D)** Directions of increasing fibroblast, monocyte and epithelial cell proportion in the PC1-PC2 and PC2-PC3 planes. Arrow coordinates are the Spearman correlations of each proportion with the two components. Grey points are the 1,935 tumors. **(E)** Simulated tumors from the fibroblast, monocyte and epithelial cell titrations projected into the primary tumor coordinate system using the global loadings, colored by the proportion of the compartment being titrated. Steps beyond the 99th percentile of that compartment in primary tumors are drawn faintly. Simulated expression from both designs was z-scored together, as one cohort, before projection. Grey points are the primary tumors. **(F)** Simulated tumors from the donor identity design, in which composition is fixed and the patient supplying the epithelial compartment varies (24 donors, 240 mixtures), projected the same way.

An axis recovered only from pooled data could reflect how the cohorts were assembled rather than the tumors in them. Cohort explained 0.02% of the variance in PC1 and 0.34% in PC2, and platform 0.00% and 0.04%, and tumors did not separate by cohort or platform within the plane (Fig. S11A, B). Recomputing the PCA within each cohort recovered the same axes, with correlations against the global loadings of 0.394 to 0.901 for PC1, 0.469 to 0.680 for PC2 and 0.314 to 0.732 for PC3 (Fig. 4C). Global PC1 matched the first within-cohort component in two cohorts and the second to fourth in the remaining eight (Fig. S11D). To place the two manipulations of Figure 2 in this space, we projected the simulated tumors onto the global loadings, scaling both designs together as one additional cohort, as each primary cohort had been scaled before pooling. Within the compositional range primary tumors occupy, titrating fibroblast, monocyte or epithelial content spread tumors 1.83 times as far along PC1 as varying the epithelial donor at fixed composition, and 0.89 times as far along PC2 (Fig. 4E, F). Epithelial content moved tumors along both components, monocyte content along PC1 alone and fibroblast content mainly along PC2, while donor identity spread tumors 1.32 times as far along PC2 as along PC1.

### Composition predicts subtype labels and accounts for their survival association

Composition and subtype covary across primary tumors, which leaves open how much composition explains the label. Recovering the label from proportions alone, in cohorts the classifier has never seen, puts a lower bound on it. We trained multinomial classifiers on inferred proportions and evaluated them by leave-one-dataset-out cross-validation. Composition explained subtype probability variance in the order MES (adjusted R² = 0.674), IMR (0.479), DIF (0.255) and PRO (0.241), while cohort membership alone explained none (−0.003 to 0.002) (Fig. 5A, Table S6). Held-out labels were recovered at ROC-AUC 0.963 for MES, 0.919 for IMR, 0.865 for PRO and 0.817 for DIF (Fig. 5B, 5C), and no held-out cohort fell below 0.918 for MES or 0.681 for DIF. Every held-out cohort exceeded the 95th percentile of a within-cohort permutation null by at least 0.14, and random forests performed within 0.03 of the multinomial model, so the recovery is not a property of the model class (Fig. S13). Expression principal components recovered the labels more accurately (MES 0.978, IMR 0.955, DIF 0.969, PRO 0.985), as features drawn from the matrix the labels are defined on should, but adding composition to them changed performance by at most 0.008 (Fig. S13).

**Figure 5.**
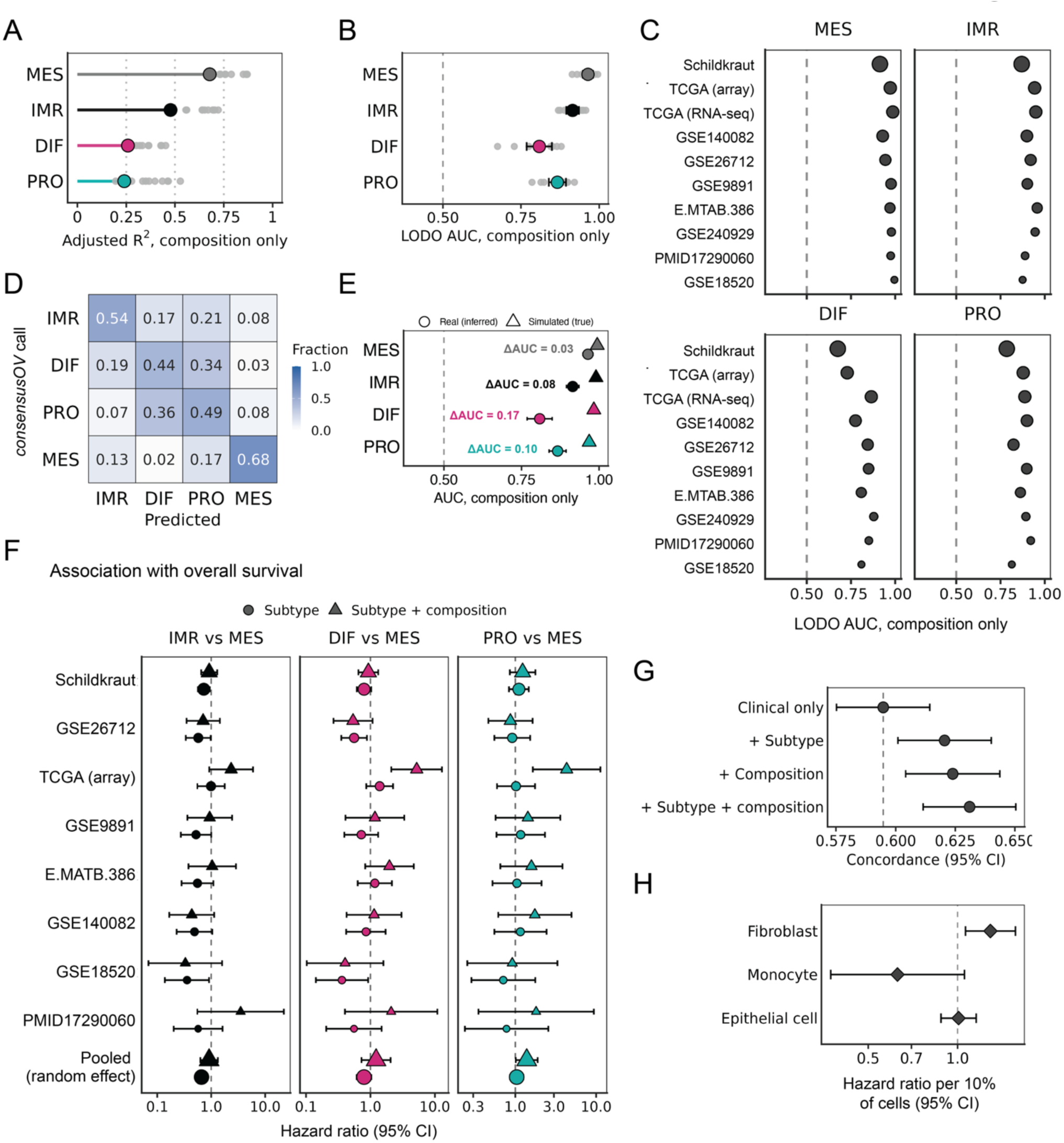
Recovery of subtype labels and survival association from composition alone. **(A)** Adjusted R² from a model of each subtype probability on the nine inferred proportions, across the 1,935 unique tumors. Grey points are the ten per-cohort values; the colored point is the value fitted across all cohorts. **(B)** Leave-one-dataset-out classification of the consensusOV call from inferred proportions alone. Grey points are the ten held-out cohorts, the colored point their mean and bars the 95% confidence interval across folds. AUC is one versus rest per subtype within each held-out cohort. **(C)** The same fold-level values, one panel per subtype, with cohorts ordered by sample size and point area proportional to the number of tumors in the held-out cohort. **(D)** Confusion matrix for the composition-only model, pooled across folds and normalized within the true class, so each row sums to one and the diagonal is per-subtype recall. **(E)** The same model fitted on simulated tumors, where proportions are set rather than inferred, against the leave-one-dataset-out values from B. Circles are real tumors with inferred composition, triangles simulated tumors with known composition; the segment and the annotation give the difference between them. Simulated tumors were evaluated by five-fold cross-validation, since they comprise one generated set rather than separate cohorts. **(F)** Cox proportional hazards estimate for each subtype against MES, within each of the eight cohorts with survival annotation and pooled by DerSimonian-Laird random-effects meta-analysis. Circles are models containing subtype and clinical covariates, triangles are the same models with the epithelial, fibroblast and monocyte proportions added. Bars span the 95% confidence interval; pooled point area is proportional to weight. The clinical covariates available in each cohort are given in Table S4. **(G)** Harrell’s concordance for nested models, pooled across the seven cohorts with at least one clinical covariate and 922 events. Bars span the 95% confidence interval. **(H)** Pooled hazard ratios for the three compartment proportions, fitted with clinical covariates and without subtype, scaled to units of 10% of cells. Correlations among the three within each cohort are given in Fig. S14D.

Recovery was uneven across the four labels, and the confusion was specific. MES was recovered with recall 0.68 and assigned to DIF in 0.03 of cases, while DIF and PRO were largely interchangeable, with 0.34 of DIF tumors assigned to PRO and 0.37 of PRO to DIF (Fig. 5D), consistent with the lower reproducibility of these two subtypes across classifiers (*33*). Two things could produce that pattern. DIF and PRO may not differ in composition, or they may differ by less than nine compartments that can be inferred from bulk expression. Separating the two requires composition that is known rather than estimated, so we refit the same model on the simulated tumors, where composition was set by construction. All four subtypes were recovered at AUC ≥ 0.959, with the largest gaps to the inferred estimates for DIF (0.981 against 0.817) and PRO (0.959 against 0.865) and the smallest for MES (0.993 against 0.963) (Fig. 5E), placing the DIF and PRO difference below the resolution of inferred proportions rather than outside composition altogether.

If the label is largely a readout of composition, the prognostic association reported for it should weaken when composition is included in the model. We fitted Cox proportional hazards models within each of the eight cohorts with recorded outcomes (1,585 tumors, 963 deaths) and pooled them by random-effects meta-analysis, fitting within cohort so that differences in treatment era and platform could not generate the association. Relative to MES, IMR was associated with longer overall survival (pooled HR 0.635, 95% CI 0.529 to 0.762), DIF showed an association in the same direction whose interval narrowly included the null (0.784, 0.611 to 1.007), and PRO showed none (0.980, 0.810 to 1.185). Heterogeneity was I² = 0% for IMR and PRO and 40.2% for DIF, and several cohorts, including TCGA (array), showed no separation by subtype (Fig. 5F). Adding inferred proportions moved the IMR estimate toward the null (0.858, 0.588 to 1.251) and left DIF and PRO indistinguishable from MES (1.138, 0.718 to 1.804; 1.234, 0.956 to 1.593, Table S7).

Two features of that design could produce the attenuation without composition being responsible for it. A single cohort could carry the pooled estimate, and a pooled hazard ratio assumes a baseline hazard the cohorts do not share. Re-estimating with each cohort omitted in turn held the pooled values between 0.581 and 0.654 for IMR, 0.736 and 0.827 for DIF, and 0.920 and 0.992 for PRO, with the IMR interval excluding the null throughout (Fig. S14A). The proportional hazards assumption failed globally in two of the eight cohort-level fits and for a single term in three more (Fig. S14C, Table S8), so we refit the comparison on the same tumors as a single model with cohort, stage and age entered as strata, which gives each cohort its own baseline hazard. No term violated the assumption in either model (global p = 0.097 and 0.182), the subtype-only estimates reproduced the pooled values to within 3% (IMR 0.632, 0.525 to 0.760; DIF 0.771, 0.649 to 0.917; PRO 0.954, 0.787 to 1.157), and adding composition again moved each toward MES (0.730, 0.573 to 0.929; 0.905, 0.710 to 1.154; 1.087, 0.847 to 1.393). IMR remained distinguishable from MES under stratification, so composition accounts for most of the subtype association rather than all (Fig. S14E).

## DISCUSSION

Our work shows that the four reproducible HGSC subtypes are regions of a compositional plane. Setting cellular composition by design in tumors reconstituted from single cells, while holding epithelial transcriptional programs fixed, reassigned subtype within the range primary tumors occupy, with fibroblast and epithelial content carrying the dependence. Varying epithelial identity at fixed composition also changed subtype assignment, but composition moved tumors nearly twice as far through expression space. Across 2,161 primary tumors from ten cohorts, composition organized the leading axes of bulk expression, recovered subtype labels in cohorts held out entirely (ROC-AUC 0.817 to 0.963), and accounted for most of the subtypes’ survival association. When composition was known by construction rather than inferred, it separated all four subtypes, including DIF and PRO, which have been less clearly tied to the microenvironment than MES and IMR.

These results build on evidence that has accumulated in HGSC since the subtypes were defined. Deconvolution of bulk cohorts has shown that the subtypes differ in composition (*22*), and single-cell studies have tied MES to fibroblast phenotypes, IMR to immune phenotypes, and DIF to myofibroblast and mesothelial phenotypes (*25*, *45*). Mixing microdissected epithelium with its paired stroma reassigned subtype at 10-30% stromal content (*28*), showing that stroma alone can alter subtype classification. Those designs could not vary each compartment independently while holding the epithelial cells fixed, or vary the epithelial cells at fixed composition, and so could not measure how much of each label composition explains or which cell types are most impactful. Varying all nine compartments also extends the dependence to PRO, the subtype least sensitive to stroma in the earlier admixtures (*28*). Both fibroblast and epithelial content moved the call through PRO, at positions that held across tumor backgrounds and a second single-cell reference. With composition known by construction, PRO was recovered at AUC 0.959 and DIF at 0.981, against 0.865 and 0.817 when composition was inferred, so their compositional differences are real but lie in compartments that a two-compartment mixture cannot vary and that nine compartments estimated from bulk only partly resolve.

The prognostic association of the subtypes follows from this dependence. Adding subtype to clinical covariates improved concordance by 0.027, and adding composition by 0.029, with overlapping intervals, so the two carry similar prognostic information beyond stage and age. Construction and survival answer different questions. Construction shows that composition changes the subtype call when the cancer cells are held fixed, which sets the direction of the dependence, whereas survival models show that once both are measured, subtype carries little prognostic information that composition does not. Schwede and colleagues reached the parallel conclusion for their gene signatures, finding that most stroma-enriched prognostic signatures lost their association with survival after adjustment for stromal content (*28*). Fitted on their own, fibroblast content was associated with worse survival, consistent with the stromal programs previously linked to outcome in HGSC (*2*, *22*, *36*, *45*, *46*), and monocyte content trended toward better survival, in the same direction as the IMR association it tracks, while epithelial fraction was not associated with survival. The prognostic signals that persist after such adjustment are tumor intrinsic. A BRCAness expression signature (*47*), which reflects homologous recombination deficiency and is not enriched in stromal genes, remained prognostic after adjustment for stroma and stage (*28*), and chromosomal instability signatures read directly from DNA have produced a biomarker predicting chemotherapy resistance from a single genomic test (*48*, *49*).

The same dependence accounts for where and when subtype calls change. Subtype probabilities are continuous and the class boundaries fall inside the compositional range tumors occupy, so 29.2% of the 2,161 primary tumors sat within 0.25 of a second subtype, where a modest shift in composition is enough to change the label. Subtype switching between primary and recurrent tumors, and poor concordance between regions of one tumor, have been read as transcriptional plasticity and regional clonal heterogeneity (*19*, *50–52*), however both patterns arise in our simulations from composition alone. Anatomical site follows the same pattern. In the AOCS cohort, 63% of tumors in the MES-equivalent subtype were sampled from extra-ovarian sites against 19% of other subtypes, and the survival association of that subtype weakened once sampling site was accounted for (*28*). Within a patient, at fixed composition and donor, the call was stable across sites in 15 of 23 patients, and where it switched it did not consistently move toward MES. This places the reported site association largely in the stromal content of extra-ovarian tissues rather than in their epithelial cells. Bulk data cannot separate compositional from cell-intrinsic change, but serial and spatial sampling with composition measured alongside would.

HGSC is not the first cancer in which bulk subtypes have resolved to composition, and the cancers that went first show what replaced them. In colorectal cancer, the mesenchymal consensus subtype was traced to stromal gene expression (*29*, *31*), and single-cell profiling later defined intrinsic epithelial subtypes alongside it, so that tumor cell state and microenvironment are classified separately (*31*, *53*). In glioblastoma, removing genes expressed by non-malignant cells refined the transcriptional subtypes and tied the mesenchymal class to macrophage and microglial content (*54*), and single-cell analysis later found the mesenchymal signature expressed more strongly in immune cells than in malignant ones (*55*). In pancreatic cancer, decomposing bulk profiles into tumor and stromal components produced separate tumor-specific and stroma-specific subtypes (*56*). In each case, the malignant compartment and the microenvironment came to be measured as separate axes, no longer read from one label. Malignant transcriptional structure is present in HGSC, as the donor and site designs show, but in bulk profiles it is mixed with, and outweighed by, composition. Single-cell and spatial profiling (*34*, *52*, *57*), and deconvolution that estimates compartment-specific expression rather than proportions alone (*18*, *58*), can measure it directly.

Several limitations apply. The simulations were pseudobulk, which differs systematically from real bulk in genes that drive classification (*43*), so the causal claim rests on mixtures even though the dependencies recur across ten bulk cohorts. Mixing was linear and drew only on cells surviving dissociation, which depletes epithelium, enriches lymphocytes, and omits cell-cell interactions. Which compartments carry a dependence also depends on the reference (*59*). The immune compartment carrying IMR also shifted between settings, from T cells in reconstituted tumors to monocytes in deconvolved primary tumors and to macrophages and NK cells under a finer reference, and deconvolution underestimated lymphoid cells, so the immune associations here are best read as associations with immune content broadly. In primary tumors, proportions and subtype probabilities derive from the same matrix, which inflates their associations, and the survival cohorts differ in treatment era, follow-up and platform. The finding that subtype added little prognostic information once composition was modeled held in a single model stratified by cohort, stage and age, but the hazard and concordance estimates compare model fit within these cohorts rather than performance validated in held-out data.

HGSC transcriptomic subtypes are a coarse readout of tumor composition. Setting composition directly, while holding epithelial cells fixed, moved the subtype call within the composition range primary tumors occupy, and the same dependencies held across ten cohorts selected for high tumor content, where composition varies least. Much of the biology and prognosis attributed to the four classes therefore belongs to the tumor microenvironment. Cell type proportions carry the prognostic information the labels do, without discarding it at class boundaries, and can be estimated from any bulk profile in which a subtype has been called.

## MATERIALS AND METHODS

### Study design

The study asked whether bulk transcriptomic subtype assignment in HGSC is determined by tumor cellular composition, and was designed in two parts. In the first, tumors were reconstituted as pseudobulk profiles from annotated single cells, so that composition was known by construction rather than inferred. Six designs varied composition, epithelial donor identity or anatomical site while holding the other factors fixed, and each profile was assigned a subtype with consensusOV. In the second, the dependencies identified in simulation were tested in 2,161 bulk profiles of primary, treatment-naive HGSC from ten cohorts, with composition inferred by deconvolution. In these tumors, composition was related to subtype probabilities, to the leading axes of expression, to subtype labels in held-out cohorts and to overall survival. No analyses were prespecified.

### Single-cell reference and cell type annotation

Single-cell RNA sequencing data were obtained from a published HGSC atlas of treatment-naive tumors (GSE180661) (*34*). A random subset of 500,000 droplets was drawn before quality control and annotation, and after filtering 337,765 annotated cells from 41 patients were retained, spanning seven anatomical supersites and 137 patient-site combinations. The published per-cell annotations were collapsed to nine analysis compartments: epithelial cell, fibroblast, endothelial cell, monocyte, T cell, B cell, plasma cell, dendritic cell and mast cell (counts in Fig. S1A). Three cells labeled “Other” were excluded. A second single-cell HGSC reference of eight treatment-naïve tumors (GSE217517) (*35*) was processed the same way for the replication analyses, with the authors’ per-cell annotations mapped onto the same nine compartments, yielding 54,083 annotated cells.

### Pseudobulk construction

Pseudobulk profiles were constructed by sampling cells of each compartment in specified proportions and summing their counts, 5,000 cells per profile unless stated otherwise. Composition in these profiles is determined directly from the annotations rather than inferred. Six designs were used (Table S1).

#### Native

Each patient’s tumor was reconstituted from all of that patient’s annotated cells at its own composition. Patients required at least 500 total and 200 epithelial cells; 39 of 41 passed (Fig. S1D).

#### Epithelial only

The same patients, rebuilt from their epithelial cells alone.

#### Titration

One compartment was stepped from 0 to 1 in 20 steps while the ratios among the remaining compartments were held fixed, with 30 replicate mixtures per step, against a background of the median composition of the primary cohorts. Each compartment is resampled independently and cannot contribute zero cells, so all proportions were floored at 0.002 and renormalized; the recorded step value is the requested fraction. Titrations were run over all nine compartments, and repeated with the background epithelial fraction set to the 25th percentile, median and 75th percentile of the primary cohorts (0.633, 0.712 and 0.783), and with GSE217517 as the reference.

#### Donor identity

Composition was held at the primary-cohort median while the patient supplying the epithelial compartment was varied, drawing on the 24 patients contributing at least 1,500 epithelial cells. Each donor’s epithelial pool was subsampled to a common depth, and a parallel run left pools at natural depth.

#### Anatomical site

Composition and patient were held fixed while the site supplying the epithelial cells was varied, for the 23 patients with two or more separately sampled sites.

#### Unconstrained

All nine compartments were varied simultaneously, 1,900 mixtures per scheme, with proportions drawn uniformly over the simplex using BuDDI (v0.1.0) (*60*) or resampled from the inferred compositions of the primary cohorts.

### Subtype assignment

Subtypes were assigned with consensusOV (v1.32.0), which integrates four published HGSC subtyping models and returns a probability for each of C2.IMM, C4.DIF, C5.PRO and C1.MES (21). We refer to these as IMR (immunoreactive), DIF (differentiated), PRO (proliferative) and MES (mesenchymal), following the names of the original TCGA description. The call is the class with the highest probability, and the call margin is that probability minus the second highest. consensusOV requires Entrez identifiers; cohort symbols were mapped with org.Hs.eg.db (v3.22.0) (*61*) and pseudobulk Ensembl identifiers with the idMap function of EnrichmentBrowser (v2.40.0) (*62*), retaining the first match for identifiers with several and excluding unmapped identifiers.

### Primary tumor cohorts

Bulk transcriptomes were assembled from ten cohorts of primary, treatment-naive HGSC, 2,161 profiles in total, 898 by RNA-sequencing and 1,263 by microarray (Table S4). TCGA contributes both platforms, and 226 of its patients were profiled on both. All 2,161 profiles are used for descriptive analyses. Wherever tumors are treated as independent observations, each of these 226 patients is represented once, by its RNA-sequencing profile, leaving 1,935 unique tumors. All cohorts were collected under protocols approved by the institutional review boards of the originating institutions, with informed consent, and the Schildkraut data were accessed under an approved dbGaP data use request.

Gene identifiers were harmonized against a master list of 23,422 genes derived from the single-cell reference, converted to HGNC symbols, excluding mitochondrial and ribosomal protein genes, MALAT1 and NEAT1. Within each cohort, duplicated symbols were resolved by retaining the row with the highest mean expression, and matrices on a log scale were returned to linear scale.

### Deconvolution

Cell type proportions were inferred with InstaPrism (v0.1.6) against their pre-made nine-compartment HGSC reference (OV_refPhi), matching the nine compartments of the simulations (*43*). A second deconvolution used an expanded reference of fourteen populations (*44*), which combines the GSE217517 cells with adipocytes from single-nucleus data (GSE176171) (*63*), and separates NK cell and ILC from T cell, macrophage from monocyte and pDC from dendritic cell, and adds adipocyte. This reference was applied to the Schildkraut cohort only, the largest single cohort, using proportions estimated by the reference’s authors (*44*), to test whether the associations held at finer resolution rather than to replace the nine-compartment estimates. Recovery was assessed by deconvolving the 5,400 titration pseudobulks, whose composition is known, and regressing inferred on true proportion per compartment.

### Principal component analysis

Each cohort’s expression matrix was log1p-transformed if on a linear scale, filtered to genes measured in at least 80% of its samples, and quantile normalized. Genes were intersected across the ten cohorts, ranked by median absolute deviation averaged across cohorts so that no single large cohort determined the feature set, and reduced to the top 5,000. Each cohort was z-scored per gene and the cohorts concatenated, giving 5,000 genes across the 1,935 unique tumors. Principal components were computed by truncated SVD (irlba v2.3.7) (*64*). Eigenvector signs are arbitrary, so each component was oriented to correlate positively with epithelial cell proportion where |ρ| ≥ 0.2, and otherwise so that its largest loading is positive.

Components were related to composition by Spearman correlation against inferred proportions, and to subtype, cohort and platform by η². Reproducibility was assessed by refitting the PCA within each cohort and correlating its loadings against the global loadings, retaining the strongest absolute match among all within-cohort components, with confidence intervals from bootstrapping over genes. Simulated tumors were projected into this coordinate system using the global loadings. The titration and donor identity pseudobulks were z-scored together, as one additional cohort, before projection, so the two designs share a scale. Spread is the standard deviation along a component of donor means or of titration step means, with titration restricted to steps within the observed range of primary tumors (zero to the 99th percentile of each compartment).

### Subtype prediction from composition

Multinomial logistic regression (nnet v7.3-20) and random forests (randomForest v4.7-1.2, 1,000 trees) were trained to predict the consensusOV call from inferred proportions in the 1,935 unique tumors and evaluated by leave-one-dataset-out cross-validation. The nine proportions sum to one, so endothelial cell was dropped as the reference category, leaving eight compositional covariates. Feature sets compared were composition alone, the first ten expression principal components, both together, and components computed separately from published cell type marker genes, from the remaining genes and from all genes; expression components were fitted on the training cohorts in each fold and projected onto the held-out cohort. Performance is one-versus-rest ROC-AUC per subtype within each held-out cohort. A ceiling analysis refit the composition-only model on simulated tumors of known composition, and a permutation null shuffled labels within each cohort over 200 permutations (Supplementary Methods).

### Survival analysis

Overall survival was analyzed in the eight cohorts with survival data (E.MTAB.386, GSE9891, GSE18520, GSE26712, GSE140082, PMID17290060, Schildkraut and TCGA array), comprising 1,585 tumors and 963 deaths. Cox proportional hazards models were fitted within each cohort with subtype as a categorical term referenced to MES and three compositional covariates, epithelial cell, fibroblast and monocyte, scaled to units of 10% of cells. Clinical covariates were included where recorded in more than 80% of a cohort’s tumors and variable within it, which excludes stage in GSE18520 and GSE26712 and age in GSE18520 and PMID17290060 (Table S4). Stage was taken from the curatedOvarianData standardization; grade was not used, as all tumors are high grade.

Three models were fitted per cohort: subtype with clinical covariates, subtype with clinical covariates and composition, and composition with clinical covariates alone. Estimates were pooled by DerSimonian-Laird random-effects meta-analysis, with heterogeneity summarized as I² and τ². Discrimination was compared by Harrell’s concordance across nested models in the seven cohorts with at least one clinical covariate (922 events). The proportional hazards assumption was tested by Schoenfeld residuals within each cohort. Where it failed, the comparison was refit on the same tumors as a single Cox model with cohort, stage and age entered as strata, so that each cohort retains its own baseline hazard. Leave-one-cohort-out and collinearity analyses are described in the Supplementary Methods.

### Statistics

Associations between proportions and subtype probabilities are reported as the coefficient of determination from ordinary least squares and as Spearman correlation. Associations with R² > 0.195, a threshold chosen to mark associations explaining roughly a fifth of the variance, are shown in bold. Per-cohort R² values are reported with 95% confidence intervals against the pooled value. Where multiple pairs were tested, p values are from the slope term and adjusted by the Benjamini-Hochberg procedure within each analysis. Paired comparisons used the Wilcoxon signed rank test and unpaired comparisons the Wilcoxon rank sum test. Analyses were performed in R 4.5.2 and Python 3.10.12.

## Data, code, and materials availability

### Bulk transcriptomes

Nine cohorts were obtained from public repositories: GSE9891, GSE18520, GSE26712, GSE140082 and GSE240929 from GEO, and E.MTAB.386, PMID17290060 and the TCGA microarray and RNA-sequencing datasets through curatedOvarianData. Accession numbers are given in Table S4. Raw counts from the Schildkraut cohort are available through dbGaP under controlled access (phs002262.v3.p1).

### Single-cell references

GSE180661 (34) and GSE217517 (35). The expanded deconvolution reference (*44*) additionally uses adipocytes from GSE176171 (*63*). The nine-compartment deconvolution reference is distributed with InstaPrism (*43*).

### Derived data

Inferred cell type proportions, subtype probability tables, pseudobulk mixtures and all figure source data are available in the project repository.

### Code

All code is publicly available at https://github.com/NDavidsonLab/hgsc-composition-subtypes and will be archived in Zenodo upon acceptance. Preparation of the primary tumor cohorts, deconvolution and subtyping are implemented as a Snakemake workflow (7.32.4) (*65*); simulation and all downstream analyses are separate scripts in the same repository. All analyses were run in a single Docker image containing R 4.5.2 and Python 3.10.12 with the package versions listed above, which will be made available on Docker Hub upon acceptance.

### Materials

This study did not generate new materials.

## AUTHOR CONTRIBUTIONS

Conceptualization: S.T., M.L., and N.R.D.

Methodology: S.T., M.L., and N.R.D.

Software: S.T., M.L., and N.R.D.

Formal Analysis: S.T., M.L., and A.I.

Data Curation: S.T., L.G., J.R.M., J.M.S., J.A.D., and N.R.D.

Writing - Original Draft: S.T.

Writing - Review and Editing: S.T., M.L., A.I., L.G., K.A.L-M., L.J.C., L.A.S., J.R.M., B.G.B., C.S.G., J.M.S., J.A.D., and N.R.D.

Visualization: S.T., M.L., and N.R.D.

Supervision: N.R.D.

Funding Acquisition: C.S.G., J.M.S., J.A.D., and N.R.D.

## ACKNOWLEDGMENTS

We would like to thank the AACES interviewers, Christine Bard, LaTonda Briggs, Whitney Franz (North Carolina), and Robin Gold (Detroit). We also thank the individuals responsible for facilitating case ascertainment across the 10 sites including Jennifer Burczyk-Brown (Alabama); Rana Bayakly and Vicki Bennett (Georgia); the Louisiana Tumor Registry; Lisa Paddock and Manisha Narang (New Jersey); Diana Slone, Yingli Wolinsky, Steven Waggoner, Anne Heugel, Nancy Fusco, Kelly Ferguson, Peter Rose, Deb Strater, Taryn Ferber, Donna White, Lynn Borzi, Eric Jenison, Nairmeen Haller, Debbie Thomas, Vivian von Gruenigen, Michele McCarroll, Joyce Neading, John Geisler, Stephanie Smiddy, David Cohn, Michele Vaughan, Luis Vaccarello, Elayna Freese, James Pavelka, Pam Plummer, William Nahhas, Ellen Cato, John Moroney, Mark Wysong, Tonia Combs, Marci Bowling, and Brandon Fletcher (Ohio); Martin Whiteside (Tennessee); and Georgina Armstrong and the Texas Registry, Cancer Epidemiology and Surveillance Branch, Department of State Health Services. We also thank the AACES investigators Anthony J. Alberg, Elisa V. Bandera, Jill Barnholtz-Sloan, Melissa Bondy, Michele L. Cote, Ellen Funkhouser, Edward Peters, Ann G. Schwartz, Paul Terry, and Patricia G. Moorman. This study would not have been possible without the efforts of the North Carolina Central Tumor Registry and all the staff of the NCOCS. We also thank Christine Lankevich for her management of the data collection for the North Carolina Study and Rex C. Bentley for reviewing the pathology in the NCOCS. We would like to thank Rex C. Bentley and Ann M. Mills for reviewing the pathology of AACES. The authors used Claude Opus 5.5 (Anthropic) to assist with copyediting portions of the manuscript text. The authors reviewed and edited all output and take full responsibility for the content of the manuscript.

## Competing Interests

L.C.P receives funding from Bristol Myers Squibb, Janssen, and Karyopharm outside the scope of this work. The remaining authors declare no potential conflicts of interest.

## FUNDING

This work was supported by the National Cancer Institute (NCI) of the National Institutes of Health (R01 CA200854 to J.M.S. and J.A.D., R01 CA237170 to C.S.G., and J.A.D., R01 CA275974 to L.A.S., R01 CA076016 to J.M.S., K99/R00 CA277580 to L.J.C., and K99/R00 HG012945-05 to N.R.D.). S.T. and N.R.D. were supported by start-up funds provided by University of Colorado Anschutz Divisions of Gynecologic Oncology and Reproductive Sciences, Department of Obstetrics & Gynecology. The research reported in this publication utilized the High-Throughput Genomics and Cancer Bioinformatics Shared Resource at Huntsman Cancer Institute at the University of Utah and was supported by the NCI of the National Institutes of Health under award number P30CA042014. The content of the manuscript is solely the responsibility of the authors and does not necessarily represent the official views of the NCI or the National Institutes of Health.

## Supplementary Materials

This PDF file includes

Supplementary Methods

Figs. S1 to S14

Legends for Tables S1 to S8

Other Supplementary Material for this manuscript includes the following

Tables S1 to S8 (Microsoft Excel format)

## Notes

### Summary of Updates

This version has been substantially revised and expanded. The title has been updated to reflect the main conclusion. The primary tumor analysis now includes 2,161 bulk profiles from ten cohorts (previously 1,834), with TCGA patients profiled on both platforms deduplicated for prediction and survival analyses. Tumors reconstituted from annotated single cells are now built both as whole tumors and from epithelial cells alone, and a second single-cell reference was added to test robustness. New simulation designs vary composition and epithelial donor identity independently, keeping all changes within the range observed in primary tumors, to compare how far each moves tumors through expression space. Subtype prediction from cell type proportions is now evaluated in cohorts held out entirely. The survival analysis was expanded to compare the prognostic information carried by subtype and by composition, including a single model stratified by cohort, stage and age. Figures, supplementary materials and the code repository have been updated accordingly.

